# Cytokine and Chemokine Receptor Profiles in Adipose Tissue Vasculature Unravel Endothelial Cell Responses in HIV

**DOI:** 10.1101/2024.03.10.584280

**Authors:** Laventa M. Obare, Stephen Priest, Anas Ismael, Mona Mashayekhi, Xiuqi Zhang, Lindsey K. Stolze, Quanhu Sheng, Zer Vue, Kit Neikirk, Heather Beasley, Curtis Gabriel, Tecla Temu, Sara Gianella, Simon Mallal, John R. Koethe, Antentor Hinton, Samuel Bailin, Celestine N. Wanjalla

## Abstract

Chronic systemic inflammation contributes to a substantially elevated risk of myocardial infarction in people living with HIV (PLWH). Endothelial cell dysfunction disrupts vascular homeostasis regulation, increasing the risk of vasoconstriction, inflammation, and thrombosis that contribute to cardiovascular disease. Our objective was to study the effects of plasma from PLWH on endothelial cell (EC) function, with the hypothesis that cytokines and chemokines are major drivers of EC activation. We first broadly phenotyped chemokine and cytokine receptor expression on arterial ECs, capillary ECs, venous ECs, and vascular smooth muscle cells (VSMCs) in adipose tissue in the subcutaneous adipose tissue of 59 PLWH using single cell transcriptomic analysis. We used CellChat to predict cell-cell interactions between ECs and other cells in the adipose tissue and Spearman correlation to measure the association between ECs and plasma cytokines. Finally, we cultured human arterial ECs (HAECs) in plasma-conditioned media from PLWH and performed bulk sequencing to study the direct effects ex-vivo. We observed that arterial and capillary ECs expressed higher interferon and tumor necrosis factor (TNF) receptors. Venous ECs had more interleukin (IL)-1R1 and ACKR1 receptors, and VSMCs had high significant IL-6R expression. CellChat predicted ligand-receptor interactions between adipose tissue immune cells as senders and capillary ECs as recipients in TNF-TNFRSF1A/B interactions. Chemokines expressed largely by capillary ECs were predicted to bind ACKR1 receptors on venous ECs. Beyond the adipose tissue, the proportion of venous ECs and VSMCs were positively plasma IL-6. In ex-vivo experiments, HAECs cultured with plasma-conditioned media from PLWH expressed transcripts that enriched for the TNF-α and reactive oxidative phosphorylation pathways. In conclusion, ECs demonstrate heterogeneity in cytokine and chemokine receptor expression. Further research is needed to fully elucidate the role of cytokines and chemokines in EC dysfunction and to develop effective therapeutic strategies.

## Introduction

The vascular endothelium is highly pleotropic and can change phenotype from an anti-inflammatory, antithrombotic, and pro-vasodilatory state to one that favors inflammation, thrombosis, and vasoconstriction (Bloom, Islam, Lesniewski, & Donato, 2023). This switch in phenotype of endothelial cells (ECS) precedes the development of atherosclerotic cardiovascular disease (CVD) and is often referred to as endothelial dysfunction (Bonetti, Lerman, & Lerman, 2003; Davignon & Ganz, 2004). Persons living with HIV (PLWH) have ∼ 1.5 to 2-fold the risk of developing CVD than the general population, which persists despite suppression of plasma viremia on antiretroviral therapy (ART) (Freiberg et al., 2013; Shah et al., 2018), but the exact mechanisms that underlie this enhanced risk remain poorly characterized. We hypothesized that circulating factors in plasma of PLWH with virologic suppression, directly act on ECs and contribute to endothelial dysfunction. Here, we studied different types of ECs present in the subcutaneous adipose tissue from PLWH with and without metabolic disease and used *in vitro* endothelial cell cultures to measure transcriptional changes due to the direct effects of plasma from PLWH.

## Methods

### Human participants

PLWH were recruited from Vanderbilt University Medical Center (VUMC). The HIV, Adipose Tissue Immunology and Metabolism Cohort included participants on ART without detectable virus (add limit of detection of the assay) for at least 12 months and no secondary source of inflammation such as other active infections or rheumatologic conditions (ClinicalTrials.gov record NCT04451980)(Bailin et al., 2023; Wanjalla et al., 2023; Wanjalla et al., 2021). The study enrolled PLWH with a broad range of cardiometabolic health by design, with approximately one third without diabetes (hemoglobin A1c (HbA1c) levels ≤ 5.7% and a fasting blood glucose (FBG) ≤ 100 mg/dL), one third with prediabetes (HbA1c 5.7% to 6.4% and/or an FBG 100 mg/dl to 125 mg/dl), and one third with diabetes (HbA1c ≥ 6.5%, FBG ≥ 126 mg/dl, or on diabetes medications). The study was approved by the VUMC Institutional Review Board and participants provided written consent. Research procedures adhered to the standards set by the U.S. Department of Health and Human Services.

### Subcutaneous adipose tissue processing

Subcutaneous adipose tissue biopsies were performed to obtain whole tissue, including intact ECs, near the umbilicus using a liposuction catheter designed for extracting viable adipocytes and stromal vascular fraction (SVF), as previously published (Bailin et al., 2023). After cleansing and filtering, the adipose tissue was processed using mechanical and enzymatic disruption (collagenase D), and the SVF was cryopreserved in liquid nitrogen (Bailin et al., 2023).

### Single cell transcriptomics analysis

Four samples were pooled together for a single reaction (with a unique hashtag antibody) and loaded onto a Chromium Single Cell 5’ assay (10x Genomics), sequenced on the NovaSeq 6000 S2 platform (Illumina), and aligned to the human genome (hg38) using STAR v2.7.2a. Cellranger count v6.0.0 was used to call cells, Souporcell was used to genetically demultiplex the samples, and ambient RNA contamination was removed using SoupX with default parameters (Heaton et al., 2020). The R Statistical Programming package Seurat V4 was used for downstream processing including cell quality filtering, count normalization, and dimension reduction techniques as previously described (Hao et al., 2023; Stuart et al., 2019). We previously generated a user-friendly website to facilitate the interaction with the scRNA sequencing from the HATIM cohort available at (http://vimrg.app.vumc.org/).

To identify cell-type specific gene markers (ECs, vascular smooth muscle cells and immune cells), we used the FindAllMarkers function implemented in Seurat for differential gene expression using Wilcoxon rank sums and included genes that were expressed in 10% or more of cells and had a log_2_ fold change 0.25 or greater. We used clusterProfiler to perform gene set enrichment analysis utilizing both Gene Ontology (GO) biological processes and the Kyoto Encyclopedia of Genes and Genomes (KEGG), to identify enriched pathways in the different endothelial and vascular smooth muscle cell populations (Wu et al., 2021). To assess compositional differences by diabetes status, we used a moderated ANOVA test on logit transformed cell proportions as implemented in propeller (Phipson et al., 2022). To assess the relationship of cell proportion with morphometric and laboratory measurements, we used a partial Spearman’s adjusted for age, sex, body mass index (BMI) and diabetes status (except for lipid and glucose measures). Individuals with less than 20 vascular cells were excluded from this analysis due to low reproducibility. DecoupleR was used for pathway and transcription factor activity inference (Badia-I-Mompel et al., 2022). For single cell metabolic pathway enrichment, we used Single Cell Pathway Analysis (SCPA) comparing the cell type of interest with all other cell types (Bibby et al., 2022).

We used the CellChat for intercellular ligand-receptor analysis including ECs, vascular smooth muscle cells, and immune cells to identify ligand and ligand receptor pairs that might mediate endothelial inflammation in PLWH. We followed the CellChat tutorial except that population size was set to ‘TRUE’ (Jin et al., 2021).

### Primary human aorta endothelial cell culture

Primary human aorta ECs (HAEC) were purchased from Lonza at an early passage (passages <4, #CC-2535) and cultured in Vascular Cell Basal Medium (ATCC # PCS-100-030) supplemented with the endothelial growth media supplements (EGM −2 # CC-4176). The HAECs were tested for mycoplasma prior to expansion and cryopreservation. 15-20% fetal bovine serum was used to maintain the cell cultures.

### Endothelial cell cultures with plasma

HAEC cells were plated on collagen coated Flexcell plates that permit uniaxial stretch (Flexcell #UF-4001C, Figure S1). The cells were grown to 90% confluency. Once confluent, the media was aspirated, and the cells were coated with 150ul of plasma for 1hr in the incubator (37 C, 5% CO_2_). Then 2 ml of media was added to each well and the cells (media control and patient plasma) and the plates were exposed to mechanical cyclic stretching to simulate normal blood pressure (5% stretch) and high blood pressure (10%). The cells were cultured in triplicates per condition, for 48hrs before cell harvesting. We aspirated the supernatants and used a cell scraper to collect the ECs that were lysed in RLT buffer or suspended in RNAprotect. One well per sample was resuspended in RIPA buffer for protein expression analysis.

### RNA extraction

HAEC cells from the stretch experiments were harvested by cell scraping and resuspended in RLT buffer (Qiagen) supplemented with β-Mercaptoethanol. RNA and DNA isolation was performed using the Qiagen All prep kit (Qiagen #80204) following the manufacturers protocol, including the DNase treatment step. In an alternative approach, HAEC cultures were stored in RNAprotect (Qiagen #76526). The cells were retrieved by centrifugation at 2,000 rotations per minute (rpm) for 5 minutes at 4 C. RNA was isolated using the Autogen XTRACT 16+ instrument cartridge Code 610, with the XTRACT Total RNA Cultured Cells Kit (XK610-72). Dnase treatment was included. RNA was eluted in RNAse free water.

### Bulk sequencing

For the HAEC endothelial stretch, RNA extracted from the cells and underwent a quality assessment using the Agilent Bioanalyzer and the quantities measured using an RNA Qubit assay. We then produced double-stranded cDNA, which underwent SPIA amplification with the Ovation® RNA Seq System V2 kit (Tecan; P/N 7102-A01). We fragment cDNA using he Covaris LE220, and then constructed the library using NEBNext® Ultra™ II for DNA kit (NEB; P/N E7645L). We used the Agilent Bioanalyzer to assess the quality of the library and a qPCR approach with the KAPA Library Quantification Kit (Roche; P/N: KK4873) on the QuantStudio 12K device to quantify the library. We pooled the prepared libraries in equal ratios, cluster generation was performed on the NovaSeq 6000 System as per standard protocols. The NovaSeq 6000 platform executed 150 bp paired-end sequencing, aiming for 50M reads for each sample. The raw sequence outputs (FASTQ files) underwent a quality check, to evaluate read quality. The RTA and NCS (1.8.0; Illumina) managed base calling, while MultiQC (v1.7; Illumina) handled the overall data quality evaluation.

For the HAEC ECs incubated with plasma, without endothelial stretch, quality control assessment was performed as above. Library preparation was slightly different and performed using the NEBNext® rRNA Depletion kit and protocol (NEB; P/N: E7850X) per manufacturer recommendations. After ribosomal depletion, the RNA samples underwent mRNA enrichment using oligo(dT) beads, followed by fragmentation using divalent cations under elevated temperature. First-strand cDNA synthesis was performed using random hexamer primers, while second-strand cDNA synthesis was carried out using DNA Polymerase I and RNase H. End repair, A-tailing, and adapter ligation was performed to generate the final cDNA library. The rest of the steps were like those above.

### Statistical Analysis

We used Wilcoxon Rank Sums test adjusted for multiple comparisons using the Benjamini-Hochberg (BH) procedure to calculate differences in subcutaneous adipose tissue endothelial cell populations (also including vascular smooth muscle cells) by their glycemic status. The relationship between endothelial cell proportions with plasma cytokines and clinical demographics was performed using partial Spearman’s correlation adjusted for confounding variables.

Bulk RNA-seq were processed using the following procedure. Reads were trimmed to remove adapter sequences using Cutadapt v4.5) (Martin, 2011) and aligned to the Gencode GRCh38.p13 genome using STAR (v2.7.11a)(Dobin et al., 2013). Gencode v38 gene annotations were provided to STAR to improve the accuracy of mapping. Quality control on both raw reads and adaptor-trimmed reads was performed using FastQC (v0.11.9) (www.bioinformatics.babraham.ac.uk/projects/fastqc). FeatureCounts (v2.0.2)(Liao, Smyth, & Shi, 2014) was used to count the number of mapped reads to each gene. Heatmap3 (Zhao, Guo, Sheng, & Shyr, 2014) was used for cluster analysis and visualization. Significantly differential expressed genes with absolute fold change >= 2 and FDR adjusted p value <= 0.05 were detected by DESeq2 (v1.30.1) (Love, Huber, & Anders, 2014). Genome Ontology and KEGG pathway over-representation analysis was performed on differentially expressed genes using the WebGestaltR package (v0.4.6) (J. Wang, Vasaikar, Shi, Greer, & Zhang, 2017). Gene set enrichment analysis was performed using GSEA package (v4.3.2)(Subramanian et al., 2005) on database (v2022.1.Hs).

## Results

We have previously published an adipose tissue atlas with the 59 PLWH included in this study (Bailin et al., 2023). There were 20 individuals without diabetes, median age 46yrs [40-52], with prediabetes 42yrs [38,54] and with diabetes 55yrs [48,62] *p=0.05*. There was no difference in sex and there was similar exposure to thymidine analog exposure by metabolic status (*p=0.45).* Only 12% of PLWH with diabetes were on metformin, and there was a similar distribution of integrase inhibitor regimen in all three groups (*p=0.80*) (Bailin et al., 2023). In this study, We first phenotyped chemokine and cytokine receptor expression on arterial ECs, capillary ECs, venous ECs, and vascular smooth muscle cells (VSMCs) in adipose tissue in the subcutaneous adipose tissue of 59 PLWH using single cell transcriptomic analysis. We then used CellChat to predict cell-cell interactions between ECs and other cells in the adipose tissue and Spearman correlation to measure the association between ECs and plasma cytokines. Finally, we cultured human arterial ECs (HAECs) in plasma-conditioned media from PLWH and performed bulk sequencing to study the direct effects ex-vivo.

### Endothelial cell heterogeneity in cytokine receptor expression

We classified endothelial cell (EC) subsets and closely approximated pericytes and VSMCs in the SAT of 59 PLWH as shown (**Figure 1A**). The main clusters included arterial ECs, venous ECs, capillary ECs, pericytes, cycling vascular cells and VSMCs. Capillary ECs were the most abundant within adipose tissue, followed by venous ECs, pericytes and arterial ECs (**Figure 1B**). These were not statistically different in abundance by diabetes status (**Table S1**). Several cytokine-receptor transcripts were highly expressed in ECs compared to other non-immune and immune cells in the SVF, including intlerukin-3RA (IL3RA), IL-1 binding receptor 1 (IL1R1), IFNGR1, IFNAR1 and IFNAR2 (**Figure 1C, Table S2**). A similar analysis of all cytokine receptors limited to vascular cells (ECs, pericytes, and VSMCs) showed significant expression of IFNGR1 on arterial and capillary ECs compared to other vascular cells (**Figure 1D, Table S3**). Within SAT, IL1R1 was highly expressed on venous ECs, and IL-6R was highest in VSMC2 (**Figure 1D, Table S3**). IL3RA expression was highest in ECs compared to all other cells in the adipose tissue. The normalized and scaled expression of the IL3RA and IL1R1 transcripts further support expression of these receptors by ECs (**Figure 1E**). This heterogenous expression highlights the importance of understanding cytokine receptor expression, and therefore potential responsiveness, at the single cell level within the same tissue to better understand how different cell types may contribute to pathology.

**Figure 1.**
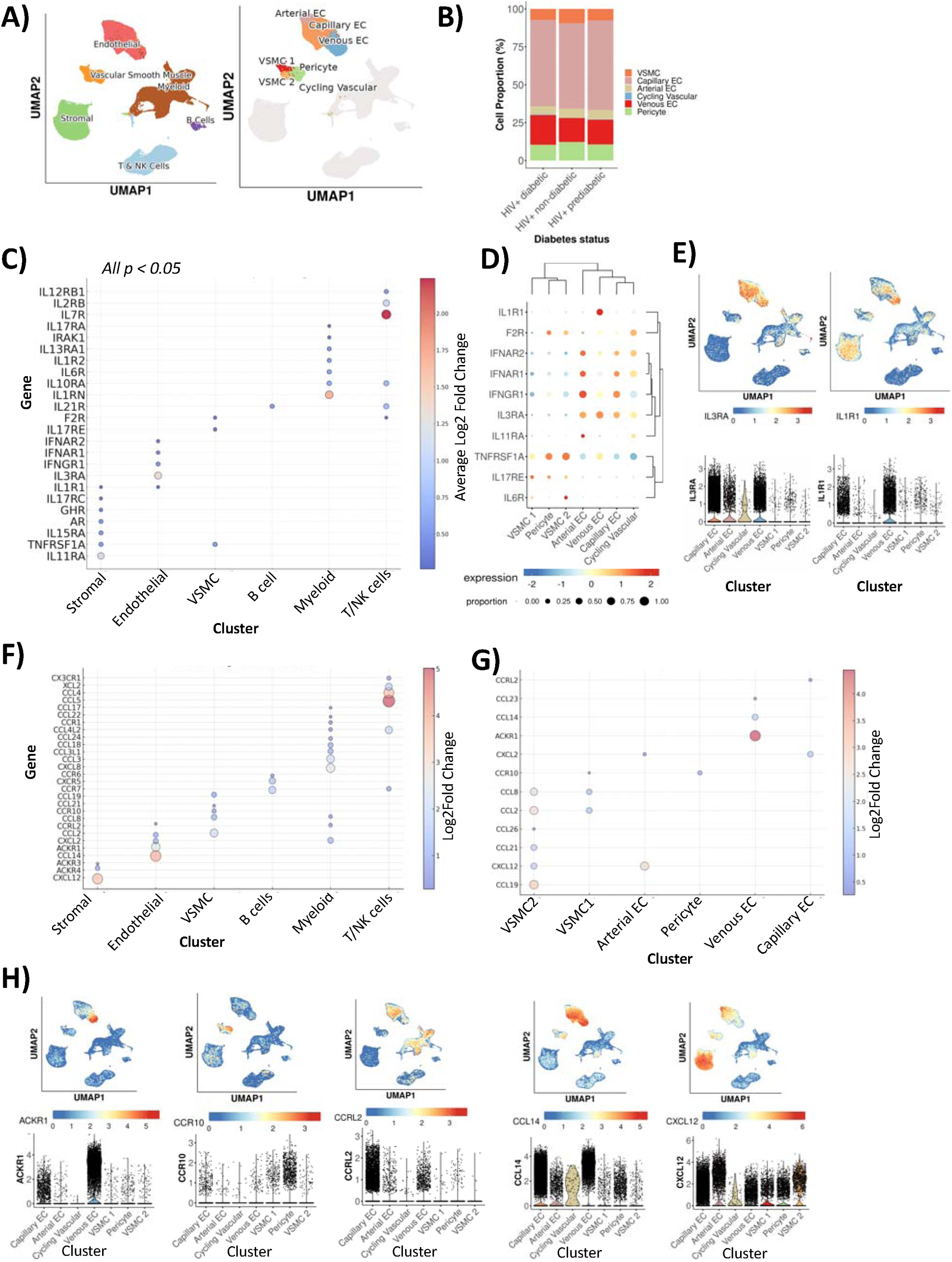
Venous, arterial, and capillary ECs have differences in cytokines receptor expression. Uniform approximation and projection (UMAP) show cell types present in adipose tissue (left) and highlighting vascular cells (A). Stacked bar plot shows each subset as a proportion of total number of vascular cells, split by metabolic status (B). Bubble plot shows cytokine receptor genes that are significantly higher in each of the cell clusters, the size of the bubble plot and color represents the average fold log2 change (see Table S2 for statistics) (color) (C). Bubble plot of select normalized and scaled cytokine receptor gene expression (y-axis) by cell type (x-axis) plotted with hierarchical clustering (D). Scaled and normalized gene expression for receptor cytokines *IL3RA,* and *IL1R1* plotted on the UMAP for all subcutaneous adipose tissue cells (top) and (E). Violin plots with major cell type (x-axis) and normalized and scaled gene expression of *IL3RA* and *IL1R1* (y-axis) (bottom) (E). The bubble plot shows differences in the expression of chemokines and chemokine receptors, with color and circle size depicting log2Fold change in all cells (F) and in vascular cells (G). ACKR1, CCR10, CCRL2, CCL14 and CXCL12 scaled and normalized expression plotted onto the UMAP (top) and violin plot by vascular cell type (bottom) (H). See Tables S2-S5

Several chemokines (CXCL2, CXCL12, CCL14, CCL22, CCL8, CCL21 and CCL19) and chemokine receptors (ACKR1, CCRL2, CCR10) were highly expressed on vascular cells compared to other cells present in the SVF (**Figure 1F, Table S4).** Notably, we also observed differences in chemokine and chemokine receptor expression in the EC subsets (**Figure 1G-H**). Arterial ECs had significant expression of CXCL2 and CXCL12 compared to other ECs. Both chemokines belong to the CXC chemokine family and have been associated with atherosclerosis (Yan, Thakur, van der Vorst, Weber, & Döring, 2021). CXCL2 is a chemokine that is more involved in acute inflammation and recruitment of monocytes (Song et al., 2023) and arterial EC-derived CXCL12 has been shown to promote atherosclerosis (Döring et al., 2019). Venous ECs, on the other hand, expressed CCL14 and CCL23 that both bind CCR1 which is involved in the recruitment of inflammatory cells to different tissues (Dyer et al., 2019). On the other hand, VSMC2 had significant expression of CCL19 and CCL21 transcripts that are both ligands for CCR7 (highly expressed on B cells, T cells and NK cells).

For chemokine receptor expression, ACKR1 was highly expressed on venous ECs and CCRL2 on capillary ECs (**Table S5**). ACKR1 acts as a scavenger that can bind and internalize several different cytokines in a manner that may regulate inflammation (Bonecchi & Graham, 2016). The differential expression of cytokine, chemokine receptors and chemokines on ECs from SAT supports the idea that plasma cytokines and chemokines may act directly on ECs and contribute to endothelial dysfunction. None of the cytokines or cytokine receptors differed by diabetes status (**Table S6**).

### Endothelial cells in SAT express both TNFRSF1A and TNFRSF1B

TNF-α is an important cytokine that has been implicated in vascular dysfunction (Zhang et al., 2009) We found several TNF receptors expressed on adipose tissue immune and non-immune cells in the SVF. TNFRSF1A was highly expressed on VSMCs, and stromal cells compared to all other cells in the SVF (**Figure 2A, Table S7**). ECs had significant expression of several TNF receptors including TNFRSF4, TNFRSF10D and CD40 when compared to other cells in the SVF. Among the vascular cells, there was significant expression of TNFRSF4 and TNFRSF1B on capillary ECs, and TNFRSF1A on VSMC2 and pericytes (**Figure 2B, Table S8)**. UMAP shows expression of TNFRSF1A and TNFRSF1B on ECs, pericytes, VSMC1 and VSMC2 (**Figure 2C**).

**Figure 2.**
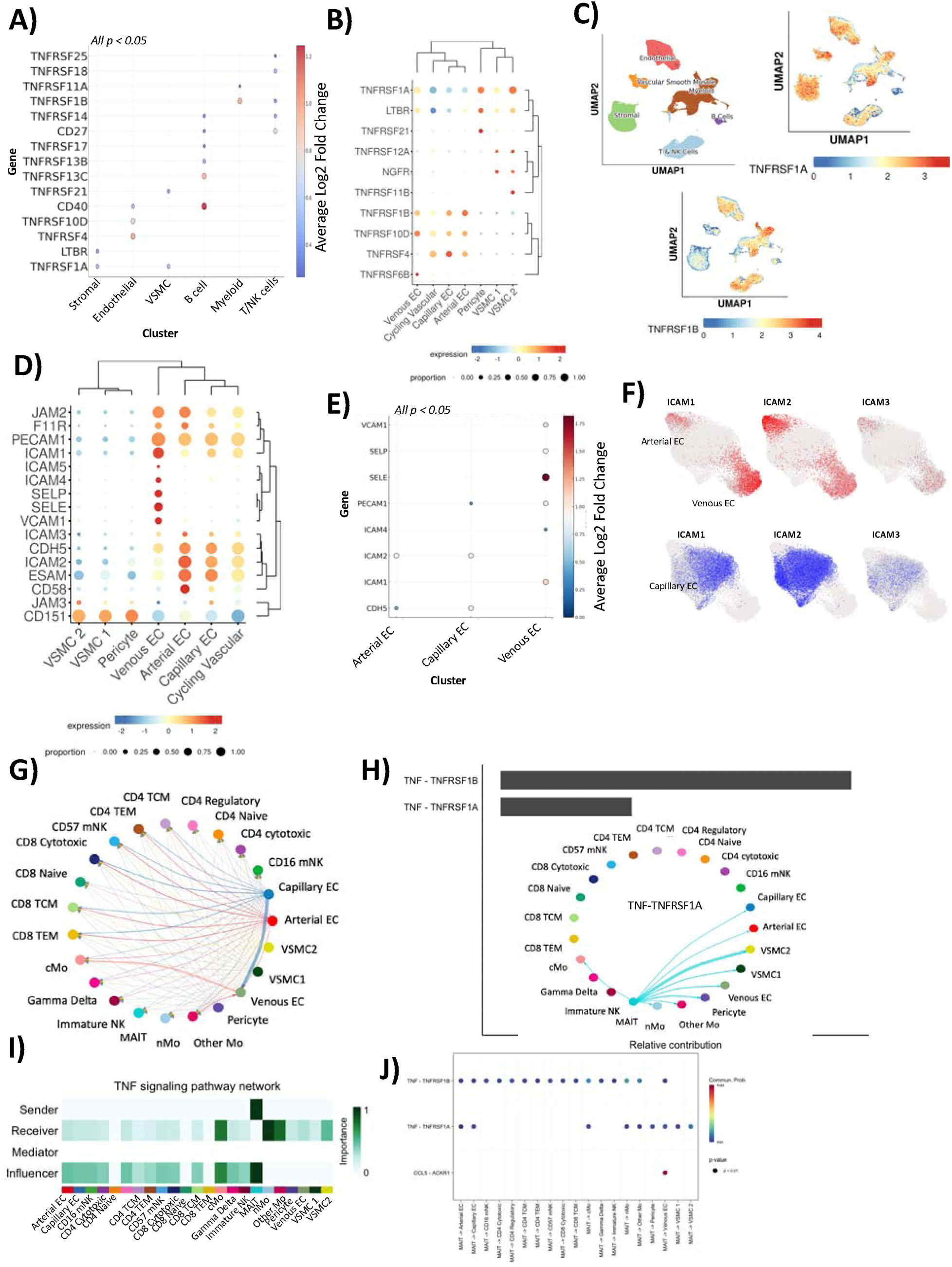
MAIT cells within SAT are predicted to be the main contributors of TNFα-that can bind TNF receptors TNFRSFR1A and TNFRSFR1B on ECs. Bubble plot for major cell types in the SVF shows differences in the expression of TNF receptors, with color and circle size depicting log2Fold change (A). Bubble plots for vascular cell populations with scaled and normalized gene expression (color) and percentage of cells expressing the gene (size). The columns and rows were plotted with hierarchical clustering for receptors and cell clusters (B). Uniform manifold approximation and projection (UMAP) with superimposed normalized and scaled expression of TNFRSFR1A and TNFRSFR1B (C). Bubble plots with expression of adhesion molecules on vascular cells with scaled and normalized gene expression (color) and percentage of cells expressing the gene (size) (D). Cell adhesion markers significantly higher in arterial, capillary, and venous ECs are depicted, with color showing fold change (E). UMAP of arterial, capillary, and venous endothelial cells with superimposed normalized and scaled gene expression of select intercellular adhesion molecules (F). Circos plot depicts communicating pairs where the line edge width represents the prevalence of each ligand-receptor interaction between endothelial cell subsets and T cells/monocytes (G). The bar plot shows the predicted ligand-receptor pairs and the relative contribution to the TNF signaling pathway network and circos plot shows interactions specific for the TNF signaling pathway network (H). The heatmap shows senders, recipients, influencers, and mediators of the TNF signaling pathway with levels of relative importance for each cell population based on the computed four network centrality measures (I). The bubble plot shows the significant ligand-receptor pairs, red dots have the highest communication probability and blue has minimum communication probability. Statistical analysis in J, one-sided permutation test. See Tables S6-S9

Inflammatory cytokines including TNF-α activate ECs leading to increased expression of cell adhesion molecules (Mackay, Loetscher, Stueber, Gehr, & Lesslauer, 1993). Cell adhesion molecules expressed on ECs are important for recruitment of leukocytes to facilitate transendothelial migration. There were apparent differences in the expression of adhesion molecules on the different ECs, VSMCs and pericytes (**Figure 2D**). We assessed cell adhesion molecules upregulated with endothelial activation in atherosclerosis (Galkina & Ley, 2007). When compared to all other cells present in SVF, we observed significant expression of intercellular cell adhesion molecule 1 (ICAM1), PECAM1, CDH5, ICAM2, SELE and SELP on ECs (**Table S9**). Among ECs, most of the adhesion molecules were expressed on venous ECs including ICAM1, VCAM1, SELE and SELP (**Figure 2E-F, Table S10).** We detected higher expression of VCAM1 on capillary ECs from diabetic compared to non-diabetic PLWH (**Table S6**). Other adhesion molecules did not differ by diabetes status.

We used CellChat to infer relationships between T cells, monocytes, and endothelial cell populations. Capillary ECs dominate the signaling networks as portrayed by the edge width in the Circos plots (**Figure 2G**). Specific analysis of the TNF signaling pathway network demonstrates that in this instance, MAIT cells were the dominant sender. The relative contribution of the TNF-TNFRSF1B pair is higher than the TNF-TNFRSF1A pair (**Figure 2H**). Arterial, capillary, and venous ECs are inferred receivers without any mediators. MAIT cells were depicted as the most important influencers in this network (**Figure 2I**). Of the different receptors, EC subsets appear to express TNFRSF1A and some TNFRSF1B (**Figure 2J**). Another predicted ligand/receptor interaction was CCL5 on MAIT cells and ACKR1 on venous ECs. Human MAIT cells can target HIV infected cells, and CCL5 is one of the main cytokines that they express upon stimulation (**Figure 2J**) (Phetsouphanh et al., 2021).

### Venous ECs are the major targets for chemokine signals

We also used CellChat to infer cell-cell communication between ECs and macrophages/fibroblasts and dendritic cells in the SVF. Most interactions were between capillary ECs and other cells (both immune and non-immune cells). The significant cytokine/chemokine signaling pathway networks included the CXCL, CCL and TNF signaling pathway networks. We observed that there was a high probability for the CXCL2 ligand and ACKR1 receptor interaction pair, which contributed the most to the CXCL signaling pathway, followed by the CXCL12-CXCR4 ligand-receptor pair (**Figure 3A**). The largest CXCL signaling was inferred to derive from the senders: capillary ECs, intermediate macrophages, and perivascular macrophages (**Figure 3B**). Other endothelial subsets also express CXCL chemokines though are deemed of lower importance based on the heatmap including arterial ECs. Venous ECs were the most important receiver, which is likely driven by the expression of the scavenger receptor ACKR1 on venous ECs. In the CellChat analysis, some cells were identified as mediators of the signaling of CXCL chemokines between the sender cells and venous ECs. These mediators included conventional dendritic cells (cDC1, cDC2B), PCOLCE+ fibroblast cells and venous ECs (autocrine). Mediators are described as cells that are likely to express proteins that can regulate the interaction between the sender and receiver. Of note, the determination of mediators and influencers is mathematical, calculated using flow betweenness to define mediators and information centrality to define influencers (Jin et al., 2021).

**Figure 3.**
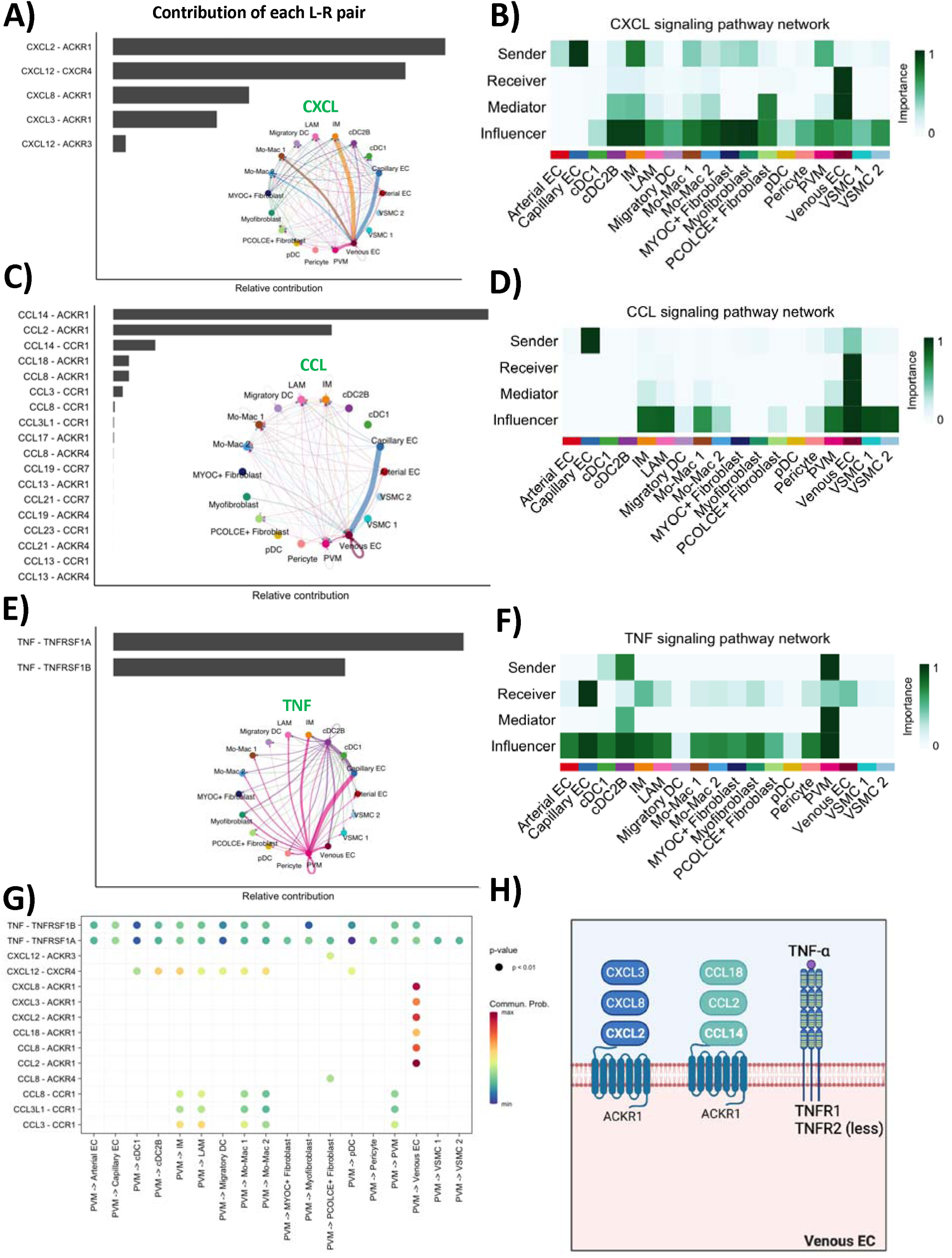
Cell-cell signaling predicted for CXCL, CCL and TNF pathways. The bar plot shows the predicted ligand-receptor pairs and the relative contribution to the CXCL signaling pathway network and the Circos plot depicts communicating pairs where the line edge width represents the prevalence of each ligand-receptor interaction (A). Heatmap shows senders, recipient, influencers, and mediators of the CXCL signaling pathway with levels of relative importance for each cell population based on the computed four network centrality measures (B). The bar plot shows the predicted ligand-receptor pairs and the relative contribution to the CCL signaling pathway network and the Circos plot depicts communicating pairs where the line edge width represents the prevalence of each ligand-receptor interaction (C). The heatmap shows senders, recipient, influencers, and mediators of the CCL signaling pathway with levels of relative importance for each cell population based on the computed four network centrality measures (D). The bar plot shows the predicted ligand-receptor pairs and the relative contribution to the TNF signaling pathway network and the Circos plot depicts communicating pairs where the line edge width represents the prevalence of each ligand-receptor interaction (E). The heatmap shows senders, recipients, influencers, and mediators of the TNF signaling pathway with levels of relative importance for each cell population based on the computed four network centrality measures (F). Bubble plot with ligand-receptor pairs of CXCL, CCL and TNF pathways (y-axis) between PVM and other cells (x-axis). The color represents the communication probability, and the size represents the p-value (G). Summary figure showing the main receptor-ligand interactions predicted for venous ECs (H).

CCL signaling between endothelial cell subsets and macrophages/fibroblasts and dendritic cells in the subcutaneous adipose tissue is also depicted (**Figure 3C**). There was a high probability for CCL14 ligand and ACKR1 receptor pair interactions, with the CCL2-ACKR1 pair contributing the most to the CCL pathway. The capillary and venous ECs were inferred to be the dominant senders of the CCL signal in this pathway, which signaled to venous ECs, and mediated by intermediate macrophages, monocyte-derived macrophages and in an autocrine manner by venous ECs (**Figure 3D**). Among the non-immune cells, pericytes and VSMCs may influence the CCL signaling pathway network, but did not turn up as mediators, despite demonstrating significant expression of CCL2.

The TNF signaling pathway network is also depicted, with TNF ligand – TNFRSF1A as the dominant ligand – receptor pair (**Figure 3E**). Of the cell types included, perivascular macrophages (PVMs) and cDC2B were the main senders and mediator of the TNF signaling pathway network, and capillary ECs were the main recipients. Venous ECs are also depicted as recipients (**Figure 3F**). TNF signaling has been shown to be important in cardiovascular disease, with direct and indirect effects on vascular function as reviewed (Zhang et al., 2009). PVMs are an important subset of macrophages that are in close proximity with ECs and can dampen or accentuate inflammatory immune responses in health and disease (Lapenna, De Palma, & Lewis, 2018). We used CellChat to look at the communication between PVM and all cells present in the SVF (**Figure 3G**). The most likely ligand-receptor communication interactions predicted were CXCL and CCL signaling pathways to venous ECs, with ACKR1 as the receptor. The communication probability of the TNF pathway was also with arterial, capillary, and venous ECs using both TNFRSF1A and TNFRSF1B (**Figure 3G**). Taken together with the assumption that CellChat identified interactions that might be meaningful, most of the interactions point to the venous ECs as major recipients within SAT. **Figure 3H** summarizes the main receptors and ligands that were predicted. ACKR1 and TNFR1 were the most predicted as contributing to the interactions. None of the chemokines or chemokine receptors differed by diabetes status (**Table S6**).

### Venous ECs are correlated with triglycerides, visceral fat volume and circulating IL-6

CellChat analysis of the SVF suggested that cytokines and chemokines within the SAT may act directly on ECs. We next explored the relationship between circulating cytokines measured in plasma (**Table S11**) and the proportion of ECs over total cells in the SVF. Arterial ECs (ρ=0.52 [0.18,0.75], p=0.004) had a positive association with fasting blood glucose levels. Venous ECs (ρ=0.41 [0.02,0.69], p=0.04) on the other hand were associated with triglycerides (**Figure 4A**). Capillary ECs were associated positively with liver mean density (ρ= 0.32 [0.40, 0.55], p=0.02) (**Figure 4B**). We also analyzed the relationships between the proportion of EC subsets and plasma cytokines adjusted for age, sex, BMI, and diabetes status. Venous ECs (ρ= 0.41 [0.14, 0.62], p=0.003) and VSMC1 (ρ= 0.30 [0.02, 0.53], p=0.04) and VSMC2 (ρ= 0.41 [0.09 0.65], p=0.01) were positively associated with plasma IL-6. Capillary ECs on the other hand were negatively associated with IL-6 (ρ= −0.36 [−0.59, −0.08], p=0.01). Finally, we observed a positive association between VSMC1 and IL-17A (ρ= 0.77 [0.31, 0.93], p=0.004) after adjusting for sex and BMI (only 19 observations) (**Figure 4C**). There was no significant relationship between the proportion of arterial ECs in the SVF and other plasma cytokines that were included in this analysis. In summary, our data suggests that within the subcutaneous adipose tissue, capillary ECs are correlated with less inflammatory cytokines while venous ECs are correlated with more inflammatory cytokines and morphometric measures.

**Figure 4.**
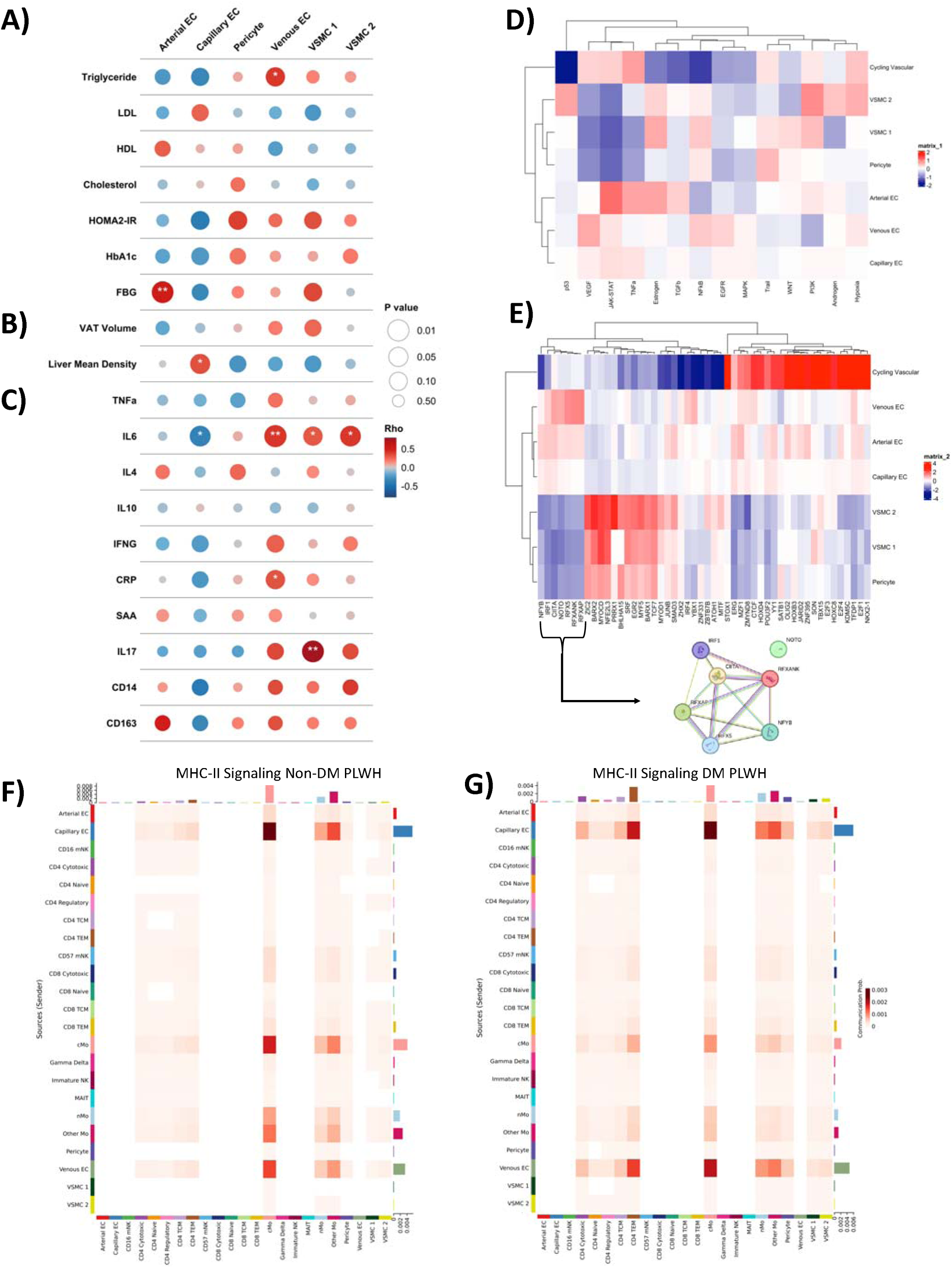
Adipose tissue capillary and venous ECs show distinct correlations with morphometric measures and plasma cytokines. Bubble plots depict the correlations between the proportion of endothelial cell subsets with laboratory measures. The size of the bubble represents the p-value, and the color is the rho for circulating glucose and lipid laboratory values (A), morphometric measures (B) and plasma cytokines (C). Statistical analysis performed using partial Spearman correlation adjusted for age, sex, BMI, and diabetes status. IL-17A analysis were adjusted for only sex and BMI due to small sample size (n=19). Heatmap shows differences in enriched pathway activity (D) and transcription factor expression of adipose tissue ECs, VSMC and pericytes (E). Heatmaps show senders (y-axis), and recipients (x-axis) of the MHCII signaling pathway with levels of relative importance for each cell population based on the computed four network centrality measures stratified by diabetes status among PLWH (F-G). See Tables S12-S13

To understand how biological functions differ between vascular cell subsets, we compared their transcriptional profiles. Several pathways came up as significant including the TNF-α signaling and the JAK-STAT pathways (**Figure 4D, Table S12**). Heatmap shows transcription factor activity was similar between arterial, venous, and capillary ECs (**Figure 4E, Table S13**). Notably, transcripts enriched in arterial, venous, and capillary ECs were overrepresented in the interferon response pathways (CIITA, IRF1, and IRF2) and TNF-α pathway (NR4A2 and IRF1). Genes that encode MHC class II and antigen presentation molecules are also expressed on ECs (**Figure S1A-C**). Functional analysis by GSEA suggests that arterial ECs are enriched for genes involved in endothelial development, antigen presentation and migration and venous ECs are enriched for genes in oxidative phosphorylation and cell adhesion (**Figure S2A-B**). Intercellular communication analysis supports the idea that arterial, capillary, and venous ECs can present antigen to monocyte-derived macrophages, lipid associated macrophages, and perivascular macrophages in both non-diabetic and diabetic PLWH (**Figure 4F**). In diabetic PLWH, arterial, capillary, and venous ECs can also signal to CD4 T cell subsets (**Figure 4G**). A comparison of metabolic pathways also showed differences between ECs (**Figure S2C, Table S14**). There are notable differences in the metabolic pathways that are overrepresented with fatty acid metabolism and triglyceride metabolism in arterial ECs and heme metabolism in venous ECs.

### Plasma from PLWH stimulates ECs with upregulation of genes that enrich for the oxidative phosphorylation and TNF-**α** pathways

To understand the direct effects of circulating factors on endothelial function, we cultured HAECs with or without plasma from PLWH with and without diabetes. We simulated normotensive stretch (**Figure 5A**), and after 24hrs-48hrs, the HAECs were lysed for RNA extraction and bulk sequencing. HAECs cultured with plasma had a fibrin like coating that developed within hours of culture. PCA plot showed co-clustering of HAECs cultured in media without versus those cultured in media with plasma from PLWH (**Figure 5B**). Differential gene expressions of ECs cultured with and without plasma are shown (**Figure 5C-D, Table S15**). We were not powered to see differential genes by diabetes status. Some of the genes that were higher in ECs cultured with plasma included FAS which is a TNF receptor and is important for antigen driven programmed cell death (**Figure 5E**). Notably, we compared FAS protein expression that we had captured (using CITE seq antibodies) in adipose tissue ECs from non-diabetic and pre-diabetic/diabetic PLWH, which also showed higher expression of FAS with diabetes (**Figure 5F)**. Using the Hallmark pathway analysis, HAECs expressed more genes that suggested activation of several pathways including MTORC1 signaling, apoptosis, oxidative phosphorylation, and TNF-α signaling via NFκB (**Figure 5G**).

**Figure 5.**
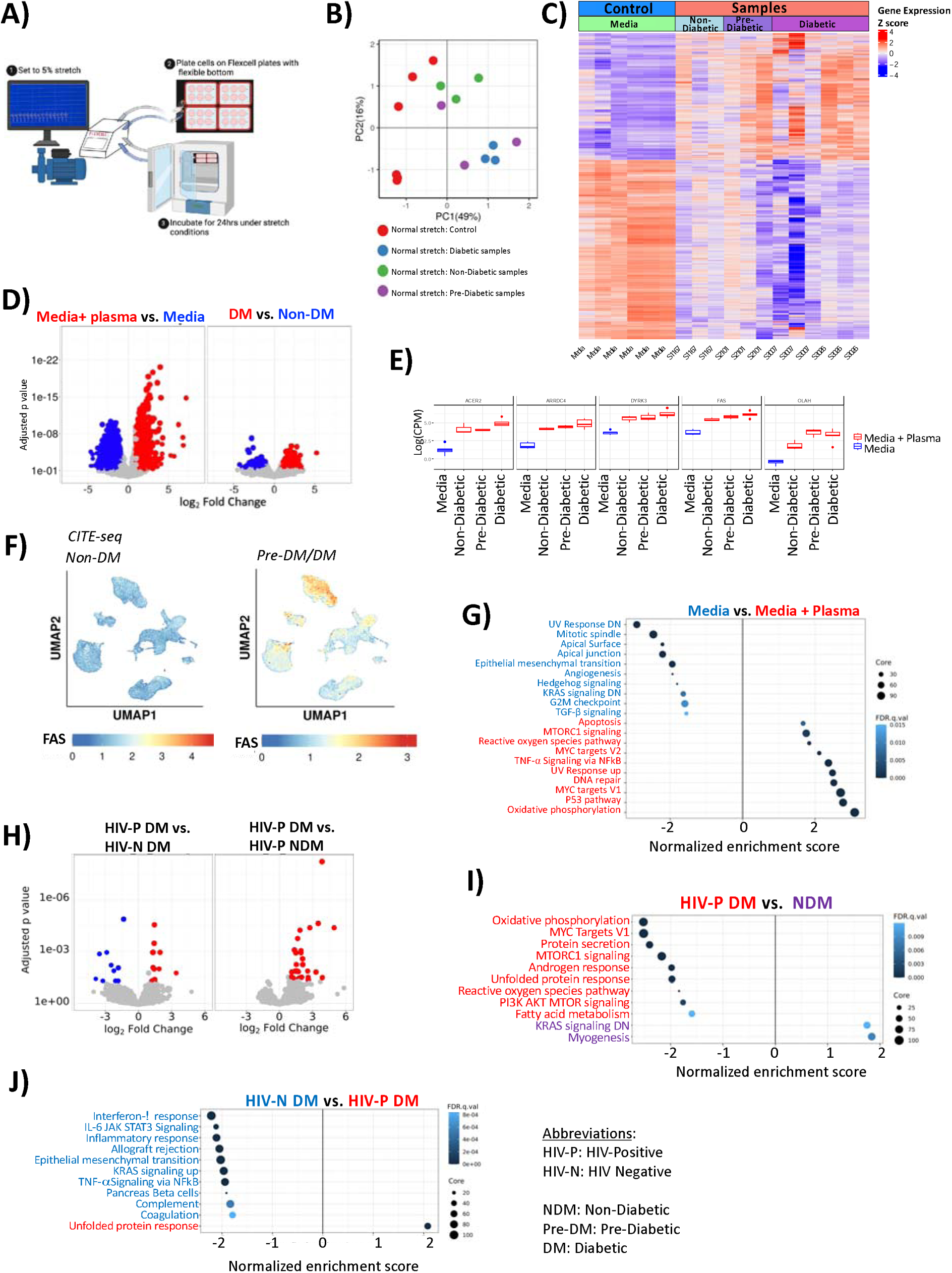
Plasma from PLWH activates HAECs to express genes that enrich several pathways including TNF-α signaling via NFKβ. Model figure showing Flexcell stretch set up with flexible plates connected to pump that allows for 5% stretch in a uni-axial plane (A). PCA plot depicting bulk RNAseq of HAECs cultured with media or plasma under normotensive or hypertensive stretch (B). Heatmap shows differential gene expression of HAECs under stretch conditions, in media only and in media supplemented with plasma (C). Volcano plots show DGE between control and plasma (left) and non-diabetic and diabetic (right) under normotensive stretch conditions (D). Box plots show the top 5 genes that differ between plasma and media samples under normotensive stretch conditions (E). UMAPs show expression of FAS protein on adipose tissue cells from non-diabetic and pre-diabetic/diabetic using the CD95 CITE-seq antibody (F). Dot plot shows the pathways in blue that are underrepresented in plasma/ normotensive stretch HAECs and those in red which are activated (G). Volcano plot shows the DGE of transcripts from HAECs cultured with media supplemented with plasma from PLWH with diabetes and HIV-negative with diabetes (left) and PLWH with and without diabetes (right) (H). Dot plot shows pathways overrepresented in HAECs in media supplemented with supernatants from PLWH with diabetes (red) compared to non-diabetic PLWH (I). Similar dot plot to compare gene transcripts between PLWH and HIV-negative with diabetes (J). See Tables S15-17

In a separate experiment, HAEC cells were cultured in non-stretch 6-well plates with plasma from PLWH with and without diabetes, as well as plasma from HIV-negative individuals with diabetes. There were differentially expressed genes (log fold change >2 and FDR < 0.05) between HAECs that had been incubated with plasma from PLWH with and without diabetes and from participants with diabetes with and without HIV (**Figure 5H, Table S16**). We observed an enriched expression of genes involved in oxidative phosphorylation, mTORC1 signaling, reactive oxygen species pathway and fatty acid metabolism in PLWH with diabetes compared to without diabetes (**Figure 5I, Table S17**). Cells that were incubated with plasma from participants with diabetes showed higher expression of genes in the interferon-γ response, IL-6, JAK/STAT3, complement and TNF-α pathway via NFκB in the HIV-negative relative to the PLWH (**Figure 5J**). This is interesting because the unfolded protein response, which was upregulated in the HIV endothelial cells, can be induced by HIV Tat protein (Fan & He, 2016).

## Discussion

In this study, we showed that interferon and tumor necrosis factor receptors were higher on arterial and capillary ECs. (IL)-1R1 and ACKR1 receptors were higher on venous ECs, and IL-6R on VSMCs. CellChat predicted TNF-TNFRSF1A/B interactions between immune cells as senders and capillary ECs as recipients. There were also predicted CXC and CCL ligands expression by capillary ECs predicted to bind ACKR1 receptors on venous ECs. The proportion of venous ECs and VSMCs were positively circulating IL-6. All ECs showed significant expression of genes that allow for MHC class II presentation of antigens to other cells. In ex-vivo experiments, HAECs cultured with plasma-conditioned media from PLWH were stimulated and transcriptionally enriched for the TNF-α and reactive oxidative phosphorylation pathways.

Adipose tissue arterial, capillary, and venous ECs in PLWH are a complex group of cells with heterogenous signaling abilities. In this study, arterial and capillary ECs were similar in their expression of IFN and TNF-α related receptors. Venous ECs on the other hand, expressed IL-1R1 and IL-6R. These differences in cytokine receptors might explain differences in their response to systemic inflammatory stimuli. IL-1 cytokine has several direct and indirect effects on ECs including the regulation of vascular permeability (Fahey & Doyle, 2019) and IL-6 is a proinflammatory cytokine that is associated with endothelial dysfunction and cardiovascular disease (Wassmann et al., 2004).

In this study, adipose tissue venous ECs and VSMCs were positively associated with plasma IL-6 while capillary ECs were negatively associated with plasma IL-6. IL-6 on ECs has long been shown to be important for the recruitment of immune cells, due to other effects on ECs including increased expression of adhesion molecules (Romano et al., 1997). Our data support the idea that adipose tissue arterial, capillary, and venous ECs are all capable of upregulating MHC Class II genes, and therefore of antigen presentation (Pober, Merola, Liu, & Manes, 2017). Furthermore, the expression of TNF receptors like TNFRSF4 (also known as OX40) on ECs that can bind to OX40 ligand on T cells, which drives proliferation of T cells (L. Liu et al., 2022) is notable. This is important in the context of PLWH because activated ECs expressing high levels of ICAM-1 and VCAM-1 are poised to engage CD4 T cells and enhance HIV infection (Card, Abrenica, McKinnon, Ball, & Su, 2022).

ACKR1 is a chemokine receptor that is highly expressed on venous ECs, red blood cells, Purkinje neurons but not on leukocytes (Bachelerie et al., 2014). Unlike other chemokine receptors, ACKR1 lacks the ability to signal downstream and may largely function as a scavenger receptor that can bind more than 20 different chemokines. ACKR1 is highly expressed on venous ECs, and in this study, was predicted to interact with several chemokines (CXCL2, CCL14 and CCL2). In inflammatory responses, ACKR1 binding to inflammatory chemokines such as CXCL2 and CCL14, may allow for the chemokines to be internalized in ECs and increase the recruitment of leukocytes past inflamed endothelium (Cambier, Gouwy, & Proost, 2023). CXCL12 was predicted to bind CXCR4 in adipose tissue, which could be homeostatic or proinflammatory. CXCL12 has been shown to act synergistically with CXCL8 to enhance migration of different subsets of cells (Cambier et al., 2023) which may include CXCR4 CD4 T cell subsets that can be infected by HIV. PLWH on ART have chronic systemic inflammation that remains elevated compared to people without HIV (Arildsen, Sørensen, Ingerslev, Østergaard, & Laursen, 2013; Hove-Skovsgaard et al., 2017; Marincowitz et al., 2019; Mosepele et al., 2018; Subramanya et al., 2019). Beyond replicating virus, HIV proteins such as Tat (Cota-Gomez et al., 2002; K. Liu et al., 2005), Nef (T. Wang et al., 2014) and Gp120 (Anand, Rachel, & Parthasarathy, 2018; Kanmogne et al., 2007; Price, Ercal, Nakaoke, & Banks, 2005) may have direct effects on immune cells leading to inflammation. The ex-vivo analysis of arterial ECs with plasma from non-diabetic, pre-diabetic and diabetic PLWH highlights the role of the TNF-α pathway in the activation of ECs. A direct comparison of the effects of plasma from PLWH on HAECs compared to media control suggests upregulation of several pathways including MTORC1 signaling and the TNF-a pathway.

In summary, we show that adipose tissue ECs, pericytes and VSMCs are heterogeneous phenotypically and functionally different as demonstrated by cytokine and chemokine receptors. Most of the cell-cell communication is predicted to be among the most abundant of these ECs in adipose (capillary and venous), that is via chemokines and the TNF pathway. Antigen presentation potential by ECs is also notable. The strength of this analysis is the cohort of 59 PLWH with adipose tissue samples, allowing us to phenotype different tissue ECs from the same study. However, it is limited by the cross-sectional nature of the study and lack of people without HIV and with and without metabolic disease. Despite the limitations, our study highlights differences in chemokine and cytokine receptors on endothelial cells, that directly react to changes in plasma biomarkers, may have a direct effect on the functional profile of the cells. Future research should aim to explore how these receptor variations on endothelial cells influence their response to specific plasma biomarkers, potentially impacting cell function and contributing to disease progression.

## Supporting information

STables

## Acknowledgements

R01 DK112262 (JRK), the Tennessee Center for AIDS Research grant P30 AI110527 (SAM), Doris Duke CSDA 2021193 (CNW), K23 HL156759 (CNW), Burroughs Wellcome Fund 1021480 (CNW), 1021868.01 (AHJ).

## Declaration of interests

The authors have no competing interests.

## Notes

### Competing Interest Statement

The authors have declared no competing interest.

## References

Anand, A. R., Rachel, G., & Parthasarathy, D. (2018). HIV Proteins and Endothelial Dysfunction: Implications in Cardiovascular Disease. Front Cardiovasc Med, 5, 185. doi:10.3389/fcvm.2018.00185

Arildsen, H., Sørensen, K. E., Ingerslev, J. M., Østergaard, L. J., & Laursen, A. L. (2013). Endothelial dysfunction, increased inflammation, and activated coagulation in HIV-infected patients improve after initiation of highly active antiretroviral therapy. HIV Med, 14(1), 1–9. doi:10.1111/j.1468-1293.2012.01027.x

Bachelerie, F., Ben-Baruch, A., Burkhardt, A. M., Combadiere, C., Farber, J. M., Graham, G. J., Zlotnik, A. (2014). International Union of Basic and Clinical Pharmacology. [corrected]. LXXXIX. Update on the extended family of chemokine receptors and introducing a new nomenclature for atypical chemokine receptors. Pharmacol Rev, 66(1), 1–79. doi:10.1124/pr.113.007724

Badia-I-Mompel, P., Vélez Santiago, J., Braunger, J., Geiss, C., Dimitrov, D., Müller-Dott, S., Saez-Rodriguez, J. (2022). decoupleR: ensemble of computational methods to infer biological activities from omics data. Bioinform Adv, 2(1), vbac016. doi:10.1093/bioadv/vbac016

Bailin, S. S., Kropski, J. A., Gangula, R. D., Hannah, L., Simmons, J. D., Mashayekhi, M., Koethe, J. R. (2023). Changes in subcutaneous white adipose tissue cellular composition and molecular programs underlie glucose intolerance in persons with HIV. Front Immunol, 14, 1152003. doi:10.3389/fimmu.2023.1152003

Bibby, J. A., Agarwal, D., Freiwald, T., Kunz, N., Merle, N. S., West, E. E., Zhang, N. R. (2022). Systematic single-cell pathway analysis to characterize early T cell activation. Cell Rep, 41(8), 111697. doi:10.1016/j.celrep.2022.111697

Bloom, S. I., Islam, M. T., Lesniewski, L. A., & Donato, A. J. (2023). Mechanisms and consequences of endothelial cell senescence. Nat Rev Cardiol, 20(1), 38–51. doi:10.1038/s41569-022-00739-0

Bonecchi, R., & Graham, G. J. (2016). Atypical Chemokine Receptors and Their Roles in the Resolution of the Inflammatory Response. Front Immunol, 7, 224. doi:10.3389/fimmu.2016.00224

Bonetti, P. O., Lerman, L. O., & Lerman, A. (2003). Endothelial dysfunction: a marker of atherosclerotic risk. Arterioscler Thromb Vasc Biol, 23(2), 168–175. doi:10.1161/01.atv.0000051384.43104.fc

Cambier, S., Gouwy, M., & Proost, P. (2023). The chemokines CXCL8 and CXCL12: molecular and functional properties, role in disease and efforts towards pharmacological intervention. Cell Mol Immunol, 20(3), 217–251. doi:10.1038/s41423-023-00974-6

Card, C. M., Abrenica, B., McKinnon, L. R., Ball, T. B., & Su, R. C. (2022). Endothelial Cells Promote Productive HIV Infection of Resting CD4. AIDS Res Hum Retroviruses, 38(2), 111–126. doi:10.1089/AID.2021.0034

Cota-Gomez, A., Flores, N. C., Cruz, C., Casullo, A., Aw, T. Y., Ichikawa, H., Flores, S. C. (2002). The human immunodeficiency virus-1 Tat protein activates human umbilical vein endothelial cell E-selectin expression via an NF-kappa B-dependent mechanism. J Biol Chem, 277(17), 14390–14399. doi:10.1074/jbc.M108591200

Davignon, J., & Ganz, P. (2004). Role of endothelial dysfunction in atherosclerosis. Circulation, 109(23 Suppl 1), III27–32. doi:10.1161/01.CIR.0000131515.03336.f8

Dobin, A., Davis, C. A., Schlesinger, F., Drenkow, J., Zaleski, C., Jha, S., Gingeras, T. R. (2013). STAR: ultrafast universal RNA-seq aligner. Bioinformatics, 29(1), 15–21. doi:10.1093/bioinformatics/bts635

Donato, A. J., Walker, A. E., Magerko, K. A., Bramwell, R. C., Black, A. D., Henson, G. D., Seals, D. R. (2013). Life-long caloric restriction reduces oxidative stress and preserves nitric oxide bioavailability and function in arteries of old mice. Aging Cell, 12(5), 772–783. doi:10.1111/acel.12103

Dyer, D. P., Medina-Ruiz, L., Bartolini, R., Schuette, F., Hughes, C. E., Pallas, K., Graham, G. J. (2019). Chemokine Receptor Redundancy and Specificity Are Context Dependent. Immunity, 50(2), 378–389.e375. doi:10.1016/j.immuni.2019.01.009

Döring, Y., van der Vorst, E. P. C., Duchene, J., Jansen, Y., Gencer, S., Bidzhekov, K., Weber, C. (2019). CXCL12 Derived From Endothelial Cells Promotes Atherosclerosis to Drive Coronary Artery Disease. Circulation, 139(10), 1338–1340. doi:10.1161/CIRCULATIONAHA.118.037953

Fahey, E., & Doyle, S. L. (2019). IL-1 Family Cytokine Regulation of Vascular Permeability and Angiogenesis. Front Immunol, 10, 1426. doi:10.3389/fimmu.2019.01426

Fan, Y., & He, J. J. (2016). HIV-1 Tat Induces Unfolded Protein Response and Endoplasmic Reticulum Stress in Astrocytes and Causes Neurotoxicity through Glial Fibrillary Acidic Protein (GFAP) Activation and Aggregation. J Biol Chem, 291(43), 22819–22829. doi:10.1074/jbc.M116.731828

Freiberg, M. S., Chang, C. C., Kuller, L. H., Skanderson, M., Lowy, E., Kraemer, K. L., Justice, A. C. (2013). HIV infection and the risk of acute myocardial infarction. JAMA Intern Med, 173(8), 614–622. doi:10.1001/jamainternmed.2013.3728

Galkina, E., & Ley, K. (2007). Vascular adhesion molecules in atherosclerosis. Arterioscler Thromb Vasc Biol, 27(11), 2292–2301. doi:10.1161/ATVBAHA.107.149179

Hao, Y., Stuart, T., Kowalski, M. H., Choudhary, S., Hoffman, P., Hartman, A., Satija, R. (2023). Dictionary learning for integrative, multimodal and scalable single-cell analysis. Nat Biotechnol. doi:10.1038/s41587-023-01767-y

Heaton, H., Talman, A. M., Knights, A., Imaz, M., Gaffney, D. J., Durbin, R., Lawniczak, M. K. N. (2020). Souporcell: robust clustering of single-cell RNA-seq data by genotype without reference genotypes. Nat Methods, 17(6), 615–620. doi:10.1038/s41592-020-0820-1

Hove-Skovsgaard, M., Gaardbo, J. C., Kolte, L., Winding, K., Seljeflot, I., Svardal, A., Nielsen, S. D. (2017). HIV-infected persons with type 2 diabetes show evidence of endothelial dysfunction and increased inflammation. BMC Infect Dis, 17(1), 234. doi:10.1186/s12879-017-2334-8

Jin, S., Guerrero-Juarez, C. F., Zhang, L., Chang, I., Ramos, R., Kuan, C. H., Nie, Q. (2021). Inference and analysis of cell-cell communication using CellChat. Nat Commun, 12(1), 1088. doi:10.1038/s41467-021-21246-9

Kanmogne, G. D., Schall, K., Leibhart, J., Knipe, B., Gendelman, H. E., & Persidsky, Y. (2007). HIV-1 gp120 compromises blood-brain barrier integrity and enhances monocyte migration across blood-brain barrier: implication for viral neuropathogenesis. J Cereb Blood Flow Metab, 27(1), 123–134. doi:10.1038/sj.jcbfm.9600330

Lapenna, A., De Palma, M., & Lewis, C. E. (2018). Perivascular macrophages in health and disease. Nat Rev Immunol, 18(11), 689–702. doi:10.1038/s41577-018-0056-9

Liao, Y., Smyth, G. K., & Shi, W. (2014). featureCounts: an efficient general purpose program for assigning sequence reads to genomic features. Bioinformatics, 30(7), 923–930. doi:10.1093/bioinformatics/btt656

Liu, K., Chi, D. S., Li, C., Hall, H. K., Milhorn, D. M., & Krishnaswamy, G. (2005). HIV-1 Tat protein-induced VCAM-1 expression in human pulmonary artery endothelial cells and its signaling. Am J Physiol Lung Cell Mol Physiol, 289(2), L252–260. doi:10.1152/ajplung.00200.2004

Liu, L., Wu, Y., Ye, K., Cai, M., Zhuang, G., & Wang, J. (2022). Antibody-Targeted TNFRSF Activation for Cancer Immunotherapy: The Role of FcγRIIB Cross-Linking. Front Pharmacol, 13, 924197. doi:10.3389/fphar.2022.924197

Love, M. I., Huber, W., & Anders, S. (2014). Moderated estimation of fold change and dispersion for RNA-seq data with DESeq2. Genome Biol, 15(12), 550. doi:10.1186/s13059-014-0550-8

Mackay, F., Loetscher, H., Stueber, D., Gehr, G., & Lesslauer, W. (1993). Tumor necrosis factor alpha (TNF-alpha)-induced cell adhesion to human endothelial cells is under dominant control of one TNF receptor type, TNF-R55. J Exp Med, 177(5), 1277–1286. doi:10.1084/jem.177.5.1277

Marincowitz, C., Genis, A., Goswami, N., De Boever, P., Nawrot, T. S., & Strijdom, H. (2019). Vascular endothelial dysfunction in the wake of HIV and ART. FEBS J, 286(7), 1256–1270. doi:10.1111/febs.14657

Martin, M. (2011). Cutadapt removes adapter sequences from high-throughput sequencing reads. EMBnet.journal, v. 17, n. 1([S.l.]), pp. 10–12, may 2011. doi: doi: 10.14806/ej.17.1.200.

Mosepele, M., Mohammed, T., Mupfumi, L., Moyo, S., Bennett, K., Lockman, S., Triant, V. A. (2018). HIV disease is associated with increased biomarkers of endothelial dysfunction despite viral suppression on long-term antiretroviral therapy in Botswana. Cardiovasc J Afr, 29(3), 155–161. doi:10.5830/CVJA-2018-003

Phetsouphanh, C., Phalora, P., Hackstein, C. P., Thornhill, J., Munier, C. M. L., Meyerowitz, J., Klenerman, P. (2021). Human MAIT cells respond to and suppress HIV-1. Elife, 10. doi:10.7554/eLife.50324

Phipson, B., Sim, C. B., Porrello, E. R., Hewitt, A. W., Powell, J., & Oshlack, A. (2022). propeller: testing for differences in cell type proportions in single cell data. Bioinformatics, 38(20), 4720–4726. doi:10.1093/bioinformatics/btac582

Pober, J. S., Merola, J., Liu, R., & Manes, T. D. (2017). Antigen Presentation by Vascular Cells. Front Immunol, 8, 1907. doi:10.3389/fimmu.2017.01907

Price, T. O., Ercal, N., Nakaoke, R., & Banks, W. A. (2005). HIV-1 viral proteins gp120 and Tat induce oxidative stress in brain endothelial cells. Brain Res, 1045(1-2), 57–63. doi:10.1016/j.brainres.2005.03.031

Romano, M., Sironi, M., Toniatti, C., Polentarutti, N., Fruscella, P., Ghezzi, P., Mantovani, A. (1997). Role of IL-6 and its soluble receptor in induction of chemokines and leukocyte recruitment. Immunity, 6(3), 315–325. doi:10.1016/s1074-7613(00)80334-9

Rose, M. L. (1998). Endothelial cells as antigen-presenting cells: role in human transplant rejection. Cell Mol Life Sci, 54(9), 965–978. doi:10.1007/s000180050226

Shah, A. S. V., Stelzle, D., Lee, K. K., Beck, E. J., Alam, S., Clifford, S., Mills, N. L. (2018). Global Burden of Atherosclerotic Cardiovascular Disease in People Living With HIV. Circulation, 138(11), 1100–1112. doi:10.1161/CIRCULATIONAHA.117.033369

Song, J., Farris, D., Ariza, P., Moorjani, S., Varghese, M., Blin, M., Goldstein, D. R. (2023). Age-associated adipose tissue inflammation promotes monocyte chemotaxis and enhances atherosclerosis. Aging Cell, 22(2), e13783. doi:10.1111/acel.13783

Stuart, T., Butler, A., Hoffman, P., Hafemeister, C., Papalexi, E., Mauck, W. M., Satija, R. (2019). Comprehensive Integration of Single-Cell Data. Cell, 177(7), 1888–1902.e1821. doi:10.1016/j.cell.2019.05.031

Subramanian, A., Tamayo, P., Mootha, V. K., Mukherjee, S., Ebert, B. L., Gillette, M. A., Mesirov, J. P. (2005). Gene set enrichment analysis: a knowledge-based approach for interpreting genome-wide expression profiles. Proc Natl Acad Sci U S A, 102(43), 15545–15550. doi:10.1073/pnas.0506580102

Subramanya, V., McKay, H. S., Brusca, R. M., Palella, F. J., Kingsley, L. A., Witt, M. D., Haberlen, S. A. (2019). Inflammatory biomarkers and subclinical carotid atherosclerosis in HIV-infected and HIV-uninfected men in the Multicenter AIDS Cohort Study. PLoS One, 14(4), e0214735. doi:10.1371/journal.pone.0214735

Wang, J., Vasaikar, S., Shi, Z., Greer, M., & Zhang, B. (2017). WebGestalt 2017: a more comprehensive, powerful, flexible and interactive gene set enrichment analysis toolkit. Nucleic Acids Res, 45(W1), W130–W137. doi:10.1093/nar/gkx356

Wang, T., Green, L. A., Gupta, S. K., Kim, C., Wang, L., Almodovar, S., Clauss, M. (2014). Transfer of intracellular HIV Nef to endothelium causes endothelial dysfunction. PLoS One, 9(3), e91063. doi:10.1371/journal.pone.0091063

Wanjalla, C. N., Gabriel, C. L., Fuseini, H., Bailin, S. S., Mashayekhi, M., Simmons, J., Koethe, J. R. (2023). CD4+ T cells expressing CX3CR1, GPR56, with variable CD57 are associated with cardiometabolic diseases in persons with HIV. Front Immunol, 14, 1099356. doi:10.3389/fimmu.2023.1099356

Wanjalla, C. N., Mashayekhi, M., Bailin, S., Gabriel, C. L., Meenderink, L. M., Temu, T., Koethe, J. R. (2021). Anticytomegalovirus CD4 ^+^ T Cells Are Associated With Subclinical Atherosclerosis in Persons With HIV. *Arterioscler Thromb Vasc Biol*, Apr;41(4):1459–1473., ATVBAHA120315786. doi:10.1161/ATVBAHA.120.315786

Wassmann, S., Stumpf, M., Strehlow, K., Schmid, A., Schieffer, B., Böhm, M., & Nickenig, G. (2004). Interleukin-6 induces oxidative stress and endothelial dysfunction by overexpression of the angiotensin II type 1 receptor. Circ Res, 94(4), 534–541. doi:10.1161/01.RES.0000115557.25127.8D

Wu, T., Hu, E., Xu, S., Chen, M., Guo, P., Dai, Z., … Yu, G. (2021). clusterProfiler 4.0: A universal enrichment tool for interpreting omics data. Innovation (Camb), 2(3), 100141. doi:10.1016/j.xinn.2021.100141

Yan, Y., Thakur, M., van der Vorst, E. P. C., Weber, C., & Döring, Y. (2021). Targeting the chemokine network in atherosclerosis. Atherosclerosis, 330, 95–106. doi:10.1016/j.atherosclerosis.2021.06.912

Zhang, H., Park, Y., Wu, J., Chen, X., Lee, S., Yang, J., Zhang, C. (2009). Role of TNF-alpha in vascular dysfunction. Clin Sci (Lond), 116(3), 219–230. doi:10.1042/CS20080196

Zhao, S., Guo, Y., Sheng, Q., & Shyr, Y. (2014). Advanced heat map and clustering analysis using heatmap3. Biomed Res Int, 2014, 986048. doi:10.1155/2014/986048

