## Supplementary material for "Cytokine and Chemokine Receptor Profiles in Adipose Tissue Vasculature Unravel Endothelial Cell Responses in HIV": STables

**5-7 Key words:** Endothelial dysfunction, cytokines, chemokines, TNF, IL-6, arterial, venous

**Abstract**: **300 words, unstructured**

**Table S1. Endothelial subcluster proportions by diabetes status**

| **Cluster** | **HIV+ non-diabetic** | **HIV+ prediabetic** | **HIV+ diabetic** | **FDR** |
| --- | --- | --- | --- | --- |
| Arterial EC | 6.1% | 8.6% | 6.9% | 0.27 |
| Venous EC | 11.3% | 13.3% | 18.6% | 0.27 |
| VSMC 1 | 3.0% | 4.3% | 4.1% | 0.31 |
| Capillary EC | 64.6% | 61.4% | 56.9% | 0.51 |
| Pericyte | 13.0% | 10.8% | 11.2% | 0.52 |
| Cycling Vascular Cells | 0.12% | 0.13% | 0.50% | 0.55 |
| VSMC 2 | 1.9% | 1.4% | 1.8% | 0.57 |
| Linear modelling-based method was used for testing difference in cell proportions between conditions. | | | | |

**Table S11. Plasma cytokine expression levels by diabetes status**

|  | **HIV-P DM N=39** | **HIV-P NDM N=51** | **HIV-P Pre-DM N=44** | **Test Statistic** |
| --- | --- | --- | --- | --- |
| Age, yrs. | 54 [49, 58] | 45 [36, 52] | 44 [36, 56] | **0.01** |
| Sex, Male | 0.72 ^28^⁄ _39_ | 0.80 ^41^⁄ _51_ | 0.80 ^35^⁄ _44_ | 0.5 |
| LDL, mg/DL | 90 [80, 105] | 96 [84, 120] | 110 [93, 127] | **0.03** |
| Statin use, Yes | 0.63 ^22^⁄ _35_ | 0.22 ^10^⁄ _45_ | 0.32 ^14^⁄ _44_ | **0.01** |
| A1C, % | **6.8 [6.2, 8.9]** | 5.3 [4.9, 5.4] | 5.6 [5.2, 5.9] | **0.01** |
| BMI, Kg/m^2^ | **33.8 [30.3, 39.2]** | 30.7 [28.1, 34.1] | 31.8 [29.0, 35.3] | **0.01** |
| IL-4, pg/ml | 0.041 [0.00, 0.051] | 0.041 [0.028, 0.056] | 0.049 [0.041, 0.058] | 0.05 |
| IL-10, pg/ml | 0.33 [0.23, 0.47] | 0.31 [0.26, 0.53] | 0.32 [0.25, 0.42] | 0.5 |
| IL-1β, pg/ml | 0.18 [0.16, 0.22] | 0.21 [0.16, 0.26] | 0.20 [0.16, 0.24] | 0.3 |
| IL-5, pg/ml | 0.52 [0.34, 0.73] | 0.40 [0.26, 0.53] | 0.41 [0.29, 0.61] | 0.09 |
| IL12-p70, pg/ml | 0.23 [0.19, 0.29] | 0.23 [0.17, 0.36] | 0.20 [0.17, 0.29] | 0.7 |
| IL-6, pg/ml | 1.49 [1.03 2.15] | 1.22 [0.92, 1.79] | 1.35 [0.97, 1.59] | 0.2 |
| IL-8, pg/ml | 7.1 [5.6, 9.3] | 6.6 [5.3, 10.8] | 6.8 [5.4, 8.3] | 0.8 |
| IFN-γ, pg/ml | 6.0 [4.9, 10.2] | 6.4 [4.6, 9.9] | 5.7 [4.0, 7.9] | 0.4 |
| TNF-α, pg/ml | 1.31[1.03, 1.52] | 1.25 [1.03, 1.45] | 1.22 [0.95, 1.37] | 0.4 |
| CRP, pg/ml | 4581 [2973, 12570] | 4086 [1821, 8476] | 4039 [1359, 6664] | 0.2 |
| SAA, pg/ml | 4095 [2294, 8769] | 3075 [2297, 6731] | 2676 [1639, 4673] | 0.09 |
| VCAM1, pg/ml | 582 [436, 703] | 527 [435, 634] | 523 [410, 637] | 0.2 |
| ICAM1, pg/ml | 568 [482, 719] | 582 [475, 735] | 515 [443, 644] | **0.04** |
| a b c represents the lower quartile a, the median b, and the upper quartile c for continuous variables.  Tests used: Kruskal-Wallis test; Pearson test; Wilcoxon test. | | | | |

**Figure S1. Arterial, capillary, and venous ECs express genes that are important in antigen presentation**. MHC class II gene expression (A) and costimulatory receptors (B-C) that facilitate antigen presentation by ECs.

**Figure S2**. **Transcriptional activity analysis suggests differences in EC subset function.** Bubble plots show biological functions that differ by cell type using GO GSEA (A) and KEGG GSEA (B). GSEA disease specific metabolic analysis pathways are shown in the heatmap, with the q-values for a variety of metabolic pathways across different cell types, indicating the statistical significance of each pathway's activity for each cell type. The metabolic pathways are arranged vertically, and cell types are arranged horizontally.

See Table S14
